# siRNA enrichment in Argonaute 2-depleted *Blattella germanica*

**DOI:** 10.1101/2021.02.11.430758

**Authors:** José Carlos Montañés, Carlos Rojano, Guillem Ylla, Maria Dolors Piulachs, José Luis Maestro

## Abstract

**Background:** RNA interference (RNAi) is a cellular mechanism used to fight various threats, including transposons, aberrant RNA, and some types of viruses. It relies on dsRNA detection and, through a mechanism involving Dicer-2 (Dcr-2) and Argonaute 2 (AGO2), together with a small RNA fragment (siRNA) as a complementary guide, binds to an RNA and cleaves it.

**Methods:** Using the cockroach *Blattella germanica* as a model, we examined AGO2 activity by depleting its mRNA levels using RNAi and analyzing the phenotypes produced.

**Results:** Silencing *AGO2* expression had no remarkable effect on nymphal development or reproduction. dsRNA treatment triggered an immediate and transitory increase in *AGO2* expression, independently of Dcr-2 action. In addition, we analyzed the siRNAs generated after injecting a heterologous dsRNA in control and *AGO2*-depleted animals. The results revealed that siRNAs were distributed non-uniformly along the dsRNA sequence. In *AGO2*-depleted animals, the proportion of 22 nucleotide reads was higher and accumulations of reads appeared in areas less well-represented in the controls. We also detected a slight preference for cytosine as the first nucleotide in controls to be lost in *AGO2*-depleted individuals.

**Conclusions/General significance:** The siRNAs produced from a dsRNA are heterogeneously distributed along the length of the dsRNA and this arrangement depends on the sequence. AGO2 exerts its role as nuclease on the siRNA duplexes independently of its action on the homologous mRNA. This study sheds light on an extremely useful process for reverse genetics in laboratories, in addition to the design of more effective, specific, and eco-friendly pest-control strategies.

**Highlights:**

- RNA interference is used to fight cellular threats including transposons and viruses
- Silencing AGO2 in *Blattella germanica* does not affect development or reproduction
- siRNA distribution along the dsRNA sequence is non-uniform and sequence dependent
- AGO2 depletion increased the proportion of 22 nt reads in certain areas

## 1. Introduction

RNA interference (RNAi) is a mechanism that cells use to defend themselves against certain types of aggression, both internal and external. The cells detect the presence of double-stranded RNA (dsRNA) from certain types of viruses, transposons, and endogenous dsRNA from hairpin-like RNA secondary structures or bidirectional transcription [1]. A series of enzymatic reactions are activated to degrade the dsRNA and eliminate it.

Researchers can exploit the RNAi mechanism to perform functional genomics studies. To do this, the organism under study is treated with a dsRNA containing the sequence of a given mRNA. The cellular machinery of the treated individual processes the dsRNA using its RNAi pathway and triggers the degradation of the targeted mRNA, leading to a concomitant reduction of the protein it encodes. The resulting phenotypes provide information on the functional role of the knocked down protein [2]. Additionally, RNAi methods using several delivery strategies are being implemented for pest control purposes [3]. These strategies include transgenic crops expressing dsRNA, applying dsRNA to leaves or soil, delivering dsRNA via non-pathogenic insect viruses, and feeding liposome-encapsulated dsRNA [4,5].

The RNAi mechanism starts with the activation of a ribonuclease type III called *Dicer* (*Dcr*; *Dcr-2* in the case of insects) [6]. This cleaves the dsRNA, producing duplexes of between 21 and 23 nucleotides with two nucleotides overhanging at each 3’ end. These small double-stranded RNAs are refered to as small interfering RNAs (siRNAs) [1]. The siRNA duplexes are then bound to the double-stranded RNA binding protein R2D2, which, together with Dcr-2, transfers siRNA duplexes into the RNA-induced silencing complex (RISC). There, one of the RNA strands is kept (guide strand), while the other strand (passenger strand) is cut and released [7]. Using the siRNA guide strand as a template, the RISC finds the target RNA through complementary base pairing and cleaves it, leading to its degradation [1].

The cleaving activity of both the passenger strand and complementary RNA is implemented by the core component of the RISC, a nuclease known as Argonaute (AGO). Insects have different AGO proteins. The proteins AGO3 and Piwi are related to the piRNA pathway [8]. On the other hand, AGO1 is used in the micro RNA (miRNA) pathway, whereas AGO2 is used in the siRNA pathway, although may sometimes function in the miRNA pathway, particularly in the absence of AGO1 [9]. The loading of siRNA duplexes into the RISC is of the utmost importance as this determines which of the strands becomes guide strand, allowing it to be used to degrade the target RNA, if the guide strand corresponds to the complementary sequence.

The action of AGO2 at the cellular level in insects has primarily been studied in *Drosophila melanogaster*, in which it has been shown that AGO2 binds to small RNAs (approx. 21 nt) originating from retrotransposons and long stem-loop structures from repetitive sequences in the genome [10]. In addition, AGO2 has been shown to be solely responsible for mRNA cleavage activity guided by siRNAs [11]. In the cockroach *Blattella germanica*, depleting AGO2 reduces the silencing activity of specific dsRNAs [9].

In this work, which use *B. germanica* as the experimental organism, we analyze the regulation of *AGO2* expression and its function at different levels, including insect development and reproduction, together with its role in the maintenance of siRNAs. To do this, we injected a heterologous dsRNA under different experimental conditions, and analyzed the siRNAs produced and maintained by the RNAi machinery. The heterologous dsRNA has no target mRNA in the cells, allowing us to study the cellular mechanisms of dsRNA processing dissociated from the effect of the siRNAs on the target mRNA.

## 2. Material and methods

### 2.1 Insects sampling

Specimens of *B. germanica* were obtained from a colony reared on dog food and water at 29 ± 1 °C and 60-70 % relative humidity. In the experiments to study the duration of the gonadotrophic cycle and the number of offspring, the control and treated females were coupled with males once they moulted into adults. All dissections were carried out on carbon dioxide-anaesthetized insects.

### 2.2 *dsRNA synthesis and* B. germanica *treatment*

Systemic RNAi treatments were performed as previously described [12,13]. The primers used to generate the dsRNA against *B. germanica Dcr-2* (ds*Dcr-2*) and *AGO*2 (ds*AGO2*) are already described [6,9].

Two different heterologous dsRNAs were used throughout the work. The first of them was designed against the polyhedrin of the *Autographa californica* nucleopolyhedrovirus (dsPolyh), as previously described [9]. The second one was a dsRNA against a 128 bp fragment of the vector pSTBlue-1 (Novagen) (dspSTBlue), that was synthetized using the circularized vector as a template, with the primers: 5’ GATCCAGAATTCGTG 3’ and 5’ GTTTTCCCAGTCACG 3’.

In the experiments designed for identifying the functions of AGO2 in development and reproduction, we treated freshly emerged last (sixth) instar female nymphs (N6D0) or freshly emerged adult females (AdD0) with 2 µg of ds*AGO2* or with 2 µg of dsPolyh as a negative control.

To evaluate the expression changes on the RNAi mechanism enzymes induced by a dsRNA treatment, we treated female nymphs on the 4^th^ day of the penultimate instar (N5D4) with 2 µg of a heterologous dsRNA (dsPolyh) or with the equivalent volume of water as a negative control. The mRNA levels of *AGO2* and *Dcr-2* were measured 4, 8 and 16 h later.

To check if the immediate and transient increase of *AGO2* expression observed after dsRNA treatment requires Dcr-2 activity, we treated female nymphs freshly emerged to the penultimate instar (N5D0) with 2 µg of ds*Dcr-2* or with 2 µg of dspSTBlue as a negative control. Four days later, individuals of both treatments were injected with 2 µg of dsPolyh or water, and *Dcr-2* and *AGO2* mRNA levels were quantified 8 h later.

### 2.3 B. germanica *small RNA libraries*

*B. germanica* female nymphs recently emerged into the penultimate instar (N5D0) were treated with 2 µg of dspSTBlue (Control insects) or against *AGO2* (ds*AGO2*). Five days later (N5D5), both Control and ds*AGO2* animals were treated with 2 µg of a second heterologous dsRNA (dsPolyh). Three specimens from each group were dissected three days later, corresponding to the second day of the last nymphal instar (N6D2), and the whole body of these individuals (excluding the head and gut), were used to construct and sequence the small RNA libraries. Small RNAs were extracted using the miRNeasy Mini kit (Qiagen), and libraries were prepared using the NEBNext® Multiplex Small RNA Library Prep Set for Illumina® from New England Biolabs. Three library replicates from Control and ds*AGO2* treated insects were prepared. Sequencing was carried out at the Genomic Core Facility of the Pompeu Fabra University (Barcelona), using the Illumina NextSeq platform (single strand × 50 cycles). In addition, small RNA libraries from Ylla et al. (2016), were used to analyze siRNA in ds*Dcr-1* depleted individuals.

### 2.4 RNA extraction, cDNA synthesis and quantitative real-time PCR analysis

Total RNA was extracted from the whole animals, excluding the head and the digestive tube, using the GenElute™ Mammalian total RNA kit (Sigma) and RNA concentrations were measured using a nano spectrophotometer (Nabi). One µg of total RNA was then retrotranscribed using the Transcriptor First Strand cDNA Synthesis kit (Roche) as previously described [15]. Primers used to quantify *AGO2, Dcr-2*, and *Actin-5C* used as a reference gene were described in [9]. All reactions were run in duplicate or triplicate.

Results from RNA-seq analysis were validated by qRT-PCR using as template a cDNA obtained from the retrotranscription step in the library construction. Levels of three siRNAs corresponding to three different positions in the dsRNA sequence, were measured using the sequence of the siRNA as specific forward primer and the sequence of the adaptor ligated to the 3’ ends during the libraries construction as a common reverse primer as described previously [9]. Primers used to quantify siRNAs along the dsPolyh sequence were: siRNA 1: 5’ ATCCTTTCCTGGGACCCGGCAA 3’, siRNA 2: 5’ TGTTAACGACCAAGAAGTGATG 3’ and siRNA 3: 5’CCGACTATGTACCTCATGACGT 3’. As a reference gene, we used *U6* [9]. All reactions were run in duplicate or triplicate.

### 2.5 siRNA identification

Low-quality reads and adapters were removed from the libraries using Trim Galore! V0.6.0 (Babraham Bioinformatics; http://www.bioinformatics.babraham.ac.uk/projects/trim_galore/). Next, RNA fragments longer than 17 nucleotides were selected and mapped with Bowtie2 v2.2.6 [17] to the fragment of Polyh, *AGO2* and *Dcr-1* sequence corresponding to the dsRNA used in treatments, forcing zero mismatches on the first 17 nucleotides of the read “-L 17 -N 0”. Samtools v1.9 [18] was used to calculate the read depth per nucleotide of each strand. Reads were normalized as reads per million reads mapped to *B. germanica* genome plus the sequences of the heterologous dsRNAs (dsPolyh). Read length distribution was calculated using all the reads mapped only in Polyh sequence. The GenomicFeatures software [19] was used for computing the total number of reads mapping to each strand of the injected dsRNAs. In addition, we used the R package seqLogo [20] to visualize the nucleotide proportions in 22 nucleotides length reads.

In order to check if the sorting mechanism for the guide and passenger strands depends on thermodynamic differences between nucleotide pair bonds at the 5’ends of the siRNAs duplexes, we selected the 22 nt sequences showing the highest reads numbers and matched each sequence with the sequence that would correspond to it after being processed by Dcr-2 (two nt overhanging at each 3 ‘end), whether or not present in our libraries. For the calculation of the bond energies corresponding to the 4 nt at each 5’ end we used the ViennaRNA Web Services (http://rna.tbi.univie.ac.at/) (Kerpedjiev et al., 2015; see Supplementary data).

Small RNA-seq data are publicly available at Bioproject PRJNA689390.

### 2.6 Statistics analysis

Data from developmental and reproductive parameters, and gene expressions are expressed as mean ± standard error of the mean (S.E.M.), and statistical differences between data from control and treated animals in these experiments were evaluated using Student’s *t*-test with the IBM SPSS Statistics 24 package.

## 3. Results

### 3.1 AGO2 function in development and reproduction

The possible functions of AGO2 in development and reproduction were identified through two different RNAi experiments. In the first experiment, we treated freshly emerged last instar female *B. germanica* nymphs with ds*AGO2*, or dsPolyh as a negative control. ds*AGO2*-treated females presented a significant 85 % decrease in their *AGO2* mRNA levels 4 days after treatment (Fig. 1A). However, these *AGO2*-depleted individuals had no apparent morphological anomalies after molting into adults, and the durations of neither the last nymphal instar nor the first gonadotrophic cycle were affected (Fig. 1A).

**Figure 1.**
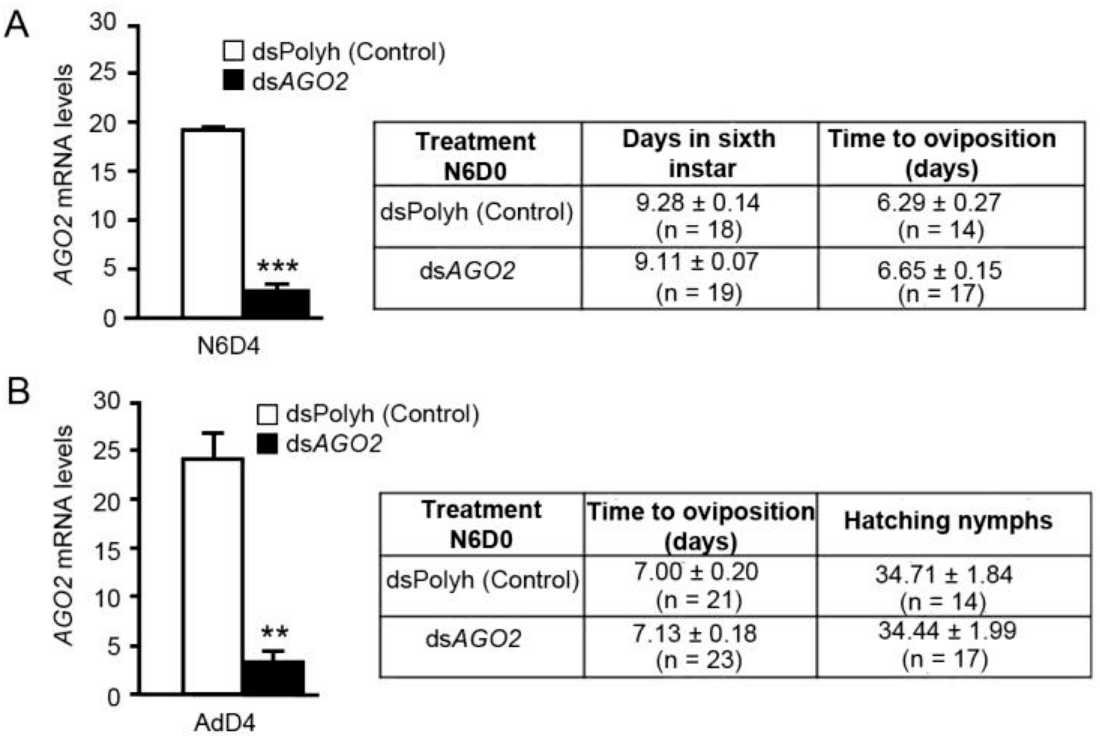
Effect of *AGO2* depletion on metamorphosis and reproduction. **A**. *AGO2* mRNA levels in 4-day-old sixth instar nymph (N6D4), treated at the day of emergence to the nymphal sixth instar with dsPolyh (Control) or ds *AGO2*. The data corresponding to the length of the sixth instar and the days that the adult females take to oviposit are indicated in annexed table. **B**. *AGO2* mRNA levels in 4-day-old adult females (AdD4), treated at the day of adult emergence with dsPolyh (Control) or ds *AGO2*. In annexed table are indicated the days that the adult females take to oviposit and the number of hatching nymphs per female in the first reproductive cycle. In A and B, data represent copies of mRNA per 1000 copies of *actin-5c* (n = 3). Data are expressed as mean ± S.E.M. Asterisks represent significant differences between Control and dsAGO2 individuals (Student’s *t*-test, **p < 0.005; ***p < 0.0001).

In a second approach, we treated freshly emerged adult females with ds*AGO2* or dsPolyh. Again, 4 days after the treatment, *AGO2* mRNA levels were 86 % lower in the treated females (Fig. 1B), but we found no differences in either the duration of the first gonadotrophic cycle, or in the number of nymphs that hatched from the ootheca formed (Fig. 1B).

### 3.2 Effect of heterologous dsRNA treatment on AGO2 and Dcr2 expression

Treating N5D4 with a heterologous dsRNA (dsPolyh) produced an immediate and transient increase in the expression of both *AGO2* (Figs. 2A) and *Dcr-2* (Fig. 2B). The *Dcr-2* mRNA increase was always more prominent than that of *AGO2* and this rise was already in evidence 4 hours after the treatment (Fig. 2B). *AGO2* mRNA levels needed 8 hours to increase 50 % increase (Figs. 2A). 16 hours after the treatment, mRNAs levels of *Dcr2* and *AGO2* were not different to their respective controls.

**Figure 2.**
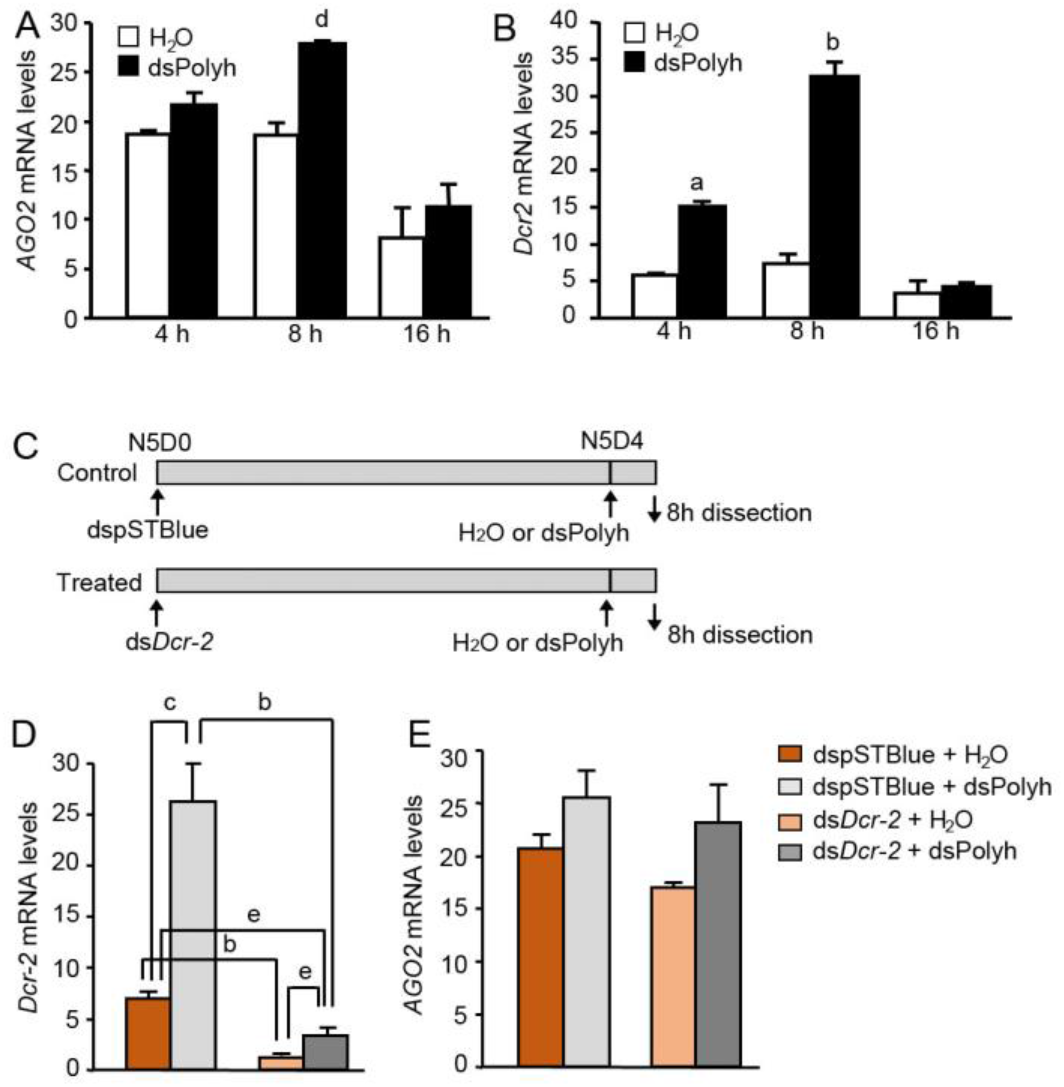
Effect of the treatment with a heterologous dsRNA on the expression of RNAi pathway enzymes. **A-B**. 4-day-old penultimate instar nymphs (N5D4) were treated with 2 µg of dsPolyh or with the equivalent volume of water. The mRNA levels of *AGO2* (A) and *Dcr-2* (B) were measured 4, 8 and 16 h later. **C-E**. Freshly emerged penultimate instar nymph (N5D0) were treated with 2 µg of ds*Dcr-2* or 2 µg of a heterologous dsRNA prepared using the pSTBlue-1 (dspSTBlue). Four days later individuals of both treatments received an injection of 2 µg of dsPolyh or the equivalent volume of water (C). *Dcr-2* (D) and *AGO2* (E) mRNA levels were measured 8 h later. Data represent copies of mRNA per 1000 copies of *actin-5c* and are expressed as the mean ± S.E.M. (n = 3-5). Statistical differences to the respective control are indicated (Student’s *t*-test, a: p<0.0001; b: p < 0.0005; c: p< 0.001; d: p< 0.005; e: p<0.05).

To check whether the observed increase in *AGO2* expression is a direct response to the dsRNA, or if *AGO2* requires Dcr2 activity to increase its expression, we treated freshly emerged N5 females with either a dsRNA against *Dcr-2* (ds*Dcr-2*), to prevent the activity of Dcr-2, or with a heterologous dsRNA (dspSTBlue, negative control) to preserve Dcr-2 activity. Four days later, individuals from both treatment groups received an injection of dsPolyh or water, and the *Dcr-2* and *AGO2* mRNA levels were quantified 8 hours later (Figure 2C, D and E). The *Dcr-2* mRNA levels had decreased in ds*Dcr-2* animals from both groups (Fig. 2D). The injection of dsPolyh increased *Dcr-2* mRNA, in both dspSTBlue and ds*Dcr-2* individuals, although much lower levels in the case of Dcr-2-depleted animals (Fig. 2D). However, *Dcr-2* mRNA levels in ds*Dcr-2* + dsPolyh subjects were lower than in dspSTBlue + H_2_O animals (Fig. 2D).

*AGO2* expression was not significantly affected by *Dcr-2* depletion, and the *AGO2* mRNA levels were similar in both dspSTBlue + H_2_O and *dsDcr-2* + H_2_O-treated females (Fig 2E). Furthermore, dsPolyh injection induced an increase in *AGO2* mRNA levels in dspSTBlue + dsPolyh (control) and ds*Dcr-2* + dsPolyh-treated individuals (23.2 % and 36.4 %, respectively); however, the differences compared with their respective controls were not statistically significant (Fig. 2E). The results therefore indicate that the increased *AGO2* expression triggered by treatment with a heterologous dsRNA is not dependent on Dcr-2 activity.

### 3.3 Polyh siRNAs analysis in Control and dsAGO2 libraries

To determine how AGO2 is implicated in the occurrence of *B. germanica* siRNAs, we analyzed the siRNAs produced in the two experimental groups: control individuals, injected sequentially with two heterologous dsRNAs (dspSTBlue + dsPolyh); and AGO2-depleted individuals, injected with ds*AGO2* + dsPolyh (Fig 3A). After assessing the *AGO2* mRNA depletion in treated individuals (77% reduction), and having found no changes in the *Dcr2* mRNA levels (Fig. S1), the small RNA libraries were prepared, sequenced, and the siRNAs produced from the dsRNA Polyh used in the treatments were analyzed (Table S1).

**Figure 3.**
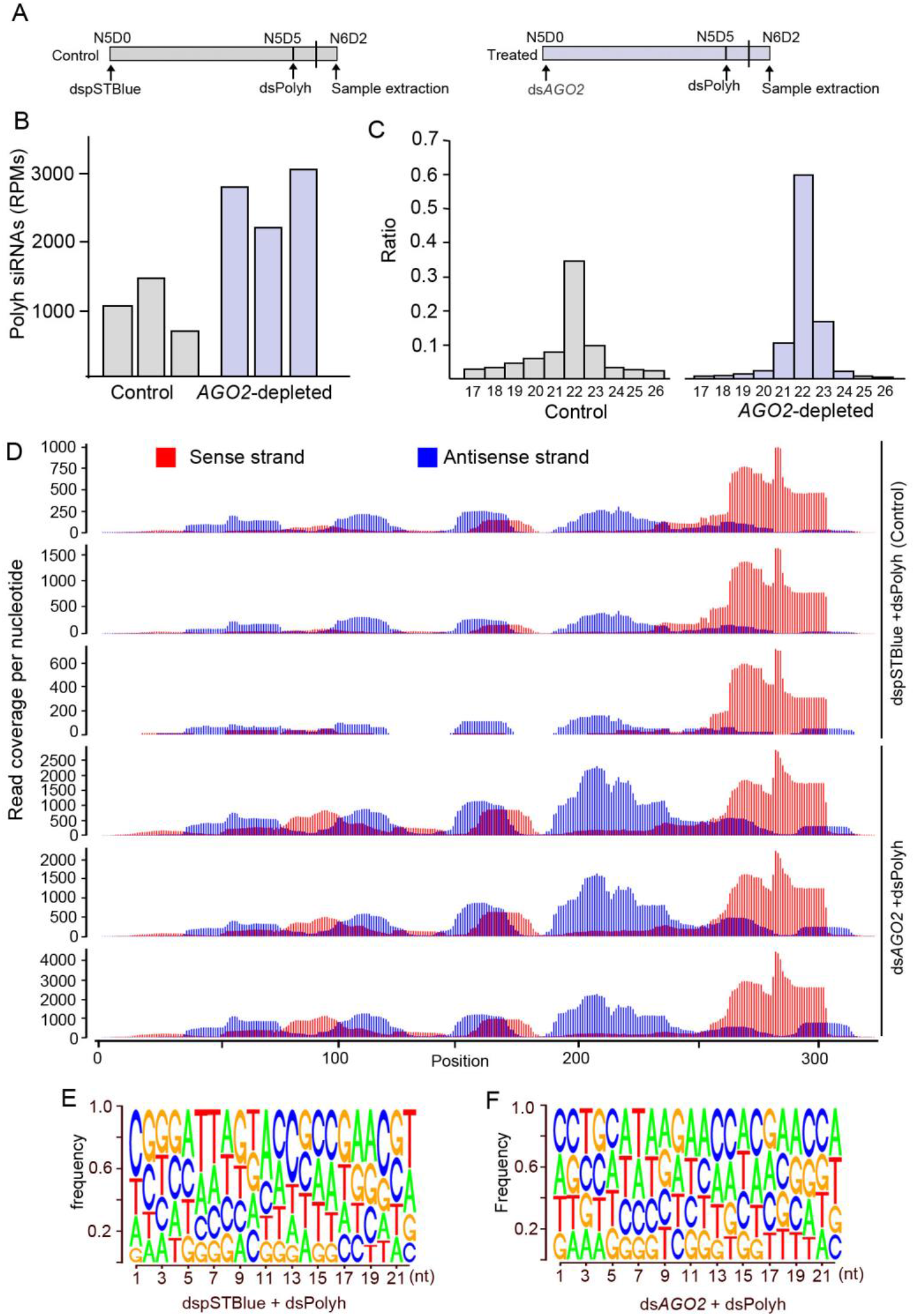
Analysis of the reads corresponding to dsRNA Polyh in Control and *AGO2*-depleted libraries. **A**. Schema explaining the experimental procedure followed to analyze the siRNAs after depletion of *AGO2* mRNA. Freshly emerged penultimate instar nymph (N5D0) were treated with 2 µg of ds*AGO2* or with 2 µg of a heterologous dspSTBlue (Control). Five days later (N5D5), both ds*AGO2* and dspSTBlue individuals were treated with 2 µg of dsPolyh. Specimens were dissected three days later, in the second day of the last nymphal instar (N6D2). **B**. Reads per million mapped reads (RPM) of Polyh siRNAs in each Control and *AGO-2* depleted library. **C**. Read length frequency of reads mapped in the dsRNA Polyh sequence. **D**. Read nucleotide coverage of the sequence of dsRNA Polyh by strand (red = sense, blue = antisense) for the 22 nt reads in Control and ds*AGO*2 libraries. **E, F**. Frequency of nucleotides in Polyh siRNAs (22 nt) reads mapping in dsRNA Polyh sequence in dspSTBlue + dsPolyh (Control) and ds*AGO2*+ dsPolyh libraries.

All the Polyh reads, from the 3 control and 3 *AGO2*-depleted libraries, were mapped individually against the sequence of dsRNA Polyh used in the second treatment (Fig 3B and Table S1). The most common size of the reads mapped to the dsPolyh sequence was 22 nucleotides (Fig 3C), and the *AGO2*-depleted libraries had a significant increase in 22 nucleotide reads, around 1.7 times that of the controls (Table S1; Fig. 3C).

The 22 nt reads produced by the treatment with the dsPolyh in both the control and treated libraries, were mapped against the dsPolyh sequence (Fig 3D). These 22 nt reads were not equally distributed along the entire length of the two RNA strands, and instead were enriched in certain areas (Fig 3D). The analyses performed mapping the reads between 17 and 51 nt gave a similar profile (Fig. S2). To validate the analysis, we repeated the experiment to construct the libraries. After small RNA extraction and retrotranscription, the relative concentrations of Polyh siRNAs were verified through quantitative real-time PCR, using oligonucleotides designed in Polyh dsRNA regions where the siRNA mapping presents high, intermediate, and low numbers of reads. The results of the qPCR confirmed the abundance differences of the different siRNAs mapped to the Polyh sequence, which had already been observed in the analysis of the libraries (Fig. S2).

Although the distribution of 22 nt Polyh reads appears similar in both the control and ds*AGO2*-treated specimens, the relative proportions of reads in some dsRNA regions differed between the two treatments (see Fig. 3D). For example, the proportion of reads mapping between nucleotides 180 and 250 on the antisense strand (blue) was higher in *AGO2*-depleted individuals than in controls. Additionally, the proportion of reads between nt 80 and 130 on the sense strand (red) was higher in *AGO2*-depleted individuals (Fig. 3D). These increased proportions point to the abundance of 22 nt reads that are not processed in *AGO2*-depleted libraries.

To detect any possible differences between the Polyh siRNA sequences from control and *AGO2*-depleted samples, we analyzed the nucleotide frequency at different positions of the 22 nt siRNA sequences (Fig 3E and F). In the Polyh siRNAs obtained from the dspSTBlue + dsPolyh (control) treatment, there was a slight preference for a cytosine in the first position at the 5’ end (Fig. 3E). This preference disappeared in the *AGO2* depleted individuals (ds*AGO2* + dsPolyh) (Fig. 3F), in which no nucleotide preference was evident at this position.

### 3.4 siRNAs from a dsRNAs targeting endogenous B. germanica RNAs

The previous results were based on an analysis of a heterologous dsRNA. We subsequently wondered if the libraries held examples of siRNAs produced by a dsRNA that targeted endogenous *B. germanica* mRNAs. We therefore analyzed the small RNA libraries, looking for siRNAs corresponding to the ds*AGO2* used in the first treatment.

In *AGO2*-depleted libraries, eight days after being treated with ds*AGO2*, it is still possible to detect a significant number of siRNA reads corresponding to this dsRNA (Table S1). These siRNAs were mapped against a *AGO2* dsRNA sequence (Fig 4A) and, in a similar way to that seen with Polyh siRNAs, the 22 nt reads of *AGO2* were not distributed homogenously along the ds*AGO2* sequence, but instead enriched certain areas of the dsRNA, with a different pattern to that observed for the siRNAs mapping to the Polyh dsRNA sequence (Fig. 4A).

**Figure 4.**
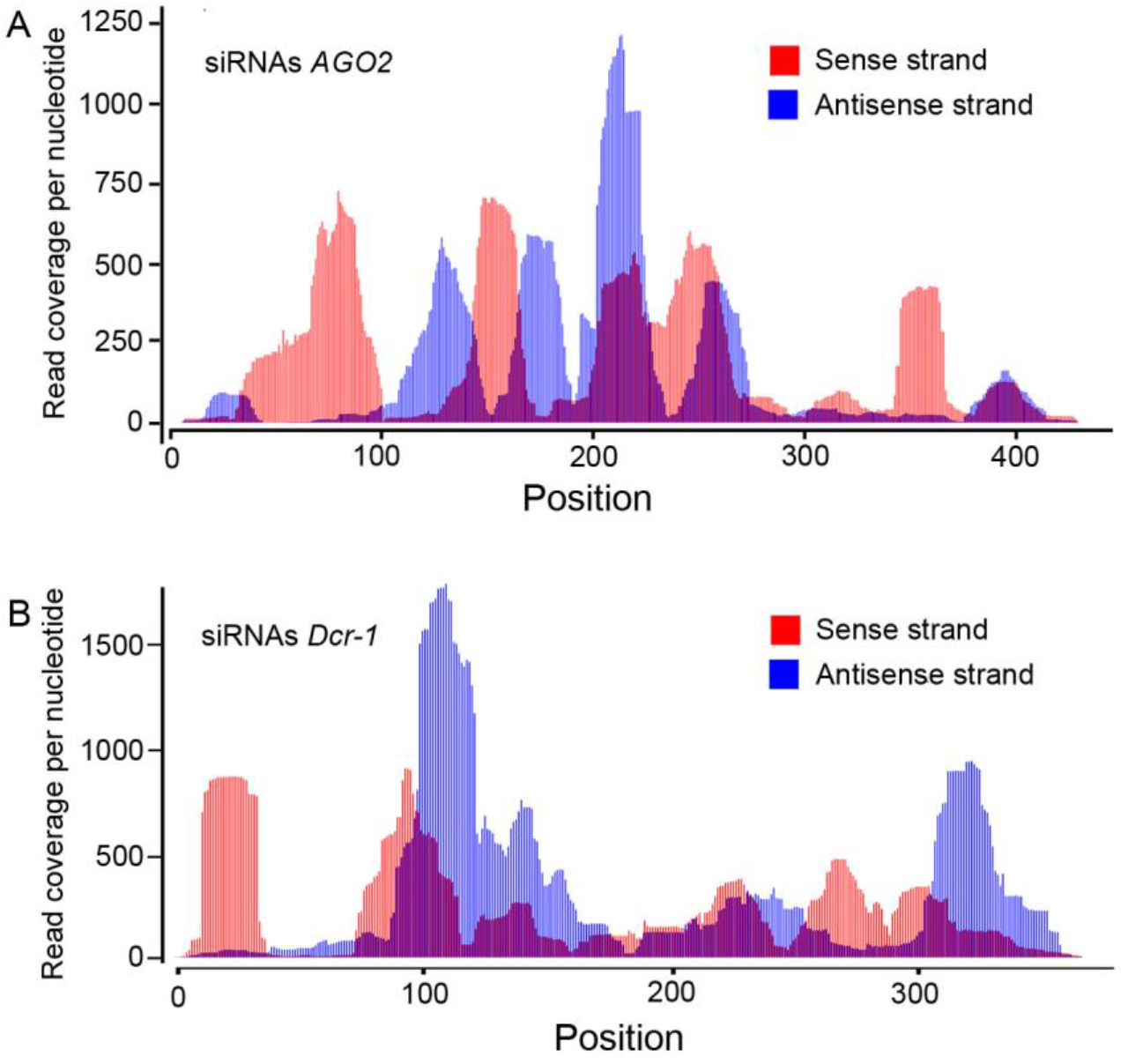
Analysis of the reads corresponding to dsRNA *AGO2* and dsRNA *Dcr-1*. **A**. Freshly emerged penultimate instar nymph (N5D0) were treated with 2 µg of ds*AGO2* or with 2 µg of a heterologous dspSTBlue (Control). Five days later (N5D5), both ds*AGO2* and dspSTBlue individuals were treated with 2 µg of dsPolyh. Specimens were dissected three days later, in the second day of the last nymphal instar (N6D2). Coverage of the 22 nt reads per nucleotide by strand (red = sense, blue = antisense) of the dsRNA *AGO2*, from 3 biological replicates together. **B**. Read coverage per nucleotide by strand (red = sense, blue = antisense) of the dsRNA *Dcr-1* for the 22 nt reads, from 2 biological replicates. These data were obtained from the libraries used in Ylla et al., (2016), using an equivalent experimental procedure.

We also analyzed the small RNA libraries produced by Ylla et al. [14], from *B. germanica* individuals treated with dsRNA against *Dcr-1*. In these libraries, we were able to identify a high number of 22 nt siRNAs mapping to the dsRNA *Dcr-1* sequence (Table S2). These 22 nt siRNAs covered the entire dsRNA sequence, with a distinct pattern that was different to those observed for Polyh and *AGO2* dsRNAs (Fig 4B). The siRNAs corresponding to the dsRNA of Polyh, used as dsMock in this experiment, showed a similar pattern to that observed in our experiments (results not shown).

### 3.5 Sorting mechanism for the guide vs the passenger strand

From the siRNA duplexes resulting from the action of Dcr-2, only one strand (the guide strand) is preserved by RISC to silence the complementary RNA. Using the 22 nt siRNAs identified in our libraries, we tested the reported model, based on *D. melanogaster*, through which the siRNA sequence retained by the RISC, the guide strand, is the one showing a thermodynamically less stable 5’ end in the siRNA duplex [7,22]. To check this model in our samples, we selected the 10-15 siRNAs with the highest number of reads for each treatment, identifying the corresponding siRNA pair on the siRNA duplex released after dsRNA cleavage by Dcr-2, in other words, the complementary sequence with 2 nt overhanging at each 3’ end, whether this sequence was in the library or not (Supplementary data). We then calculated the bond energy of the four nucleotide pairs at each end, and compared these. By checking the number of reads from each strand and the energies at each end, we determined whether the guide and passenger strand selection of the siRNA duplexes corresponding to the most abundant sequences followed the model described for *D. melanogaster*. In 62% of the duplexes analyzed (n=34) the Δ energy between the 5’ and 3’ ends of the most abundant sequence of the pair is > +0.5, a value compatible with the expected strand selection based on the calculated energies (Supplementary data). These results, do not allow us to assert that our libraries follow up the model described for *D. melanogaster*.

## 4. Discussion

The siRNA pathway, in which one of the main proteins is the nuclease AGO2, protects the organism from various threats, both internal and external, including certain types of viruses, transposons, and aberrant RNAs. Treating the last instar *B. germanica* females with a dsRNA against *AGO2* did not affect the duration of either the instar or the gonadotrophic cycle.\ Similarly, depleting AGO2 in adult females did not modify the duration of the gonadotrophic cycle or the number of offspring per female. The results indicate that AGO2 does not play a vital role in either *B. germanica* metamorphosis or female reproduction, but instead support the idea that its function involves defense against aggression and, if this is absent, the animal’s life cycle can proceed normally even when AGO2 levels are reduced. The same applies for other insect species, and similar results have been reported for *D. melanogaster*, where AGO2-mutant flies molt into the adult stage with no apparent problems, with a normal appearance and fertility [23]; and for *D. virgifera*, in which AGO2 depletion has little or no effect on larval growth, adult emergence, survival, or fecundity [24].

Treatment with a heterologous dsRNA (dsPolyh) produced an immediate and transient increase of *AGO2* and *Dcr-2* expression, which was higher in the case of *Dcr-2*. This result is similar to that obtained in *Manduca sexta* larvae treated with a dsRNA, which also produced an increase in *Dcr-2* and *AGO2* expression, again being much higher in the case of *Dcr-2* [25]. This increase in *AGO2* expression is not due to increased Dcr-2 or the occurrence of Dcr-2 products (siRNA duplexes), because *Dcr-2* depletion, in the ds*Dcr-2* treatments, did not preclude the increased *AGO2* mRNA levels triggered by the dsPolyh treatment, since similar *AGO2* mRNA levels were reached in control (dspSTBlue) and ds*Dcr-2* animals. For this reason, we presume that there is a yet unknown mechanism for sensing dsRNAs and stimulating the transcription of RNAi pathway components such as *Dcr-2* and *AGO2*. The increased *Dcr-2* and *AGO2* mRNA levels triggered by dsRNA help the cells to fight the threat generated by the dsRNA agents. In fact, *B. mori* larvae infected with cytoplasmic polyhedrosis virus, a double-stranded RNA virus, also present increased expression of both *Dcr-2* and *AGO2* [26].

In order to clarify the involvement of AGO2 in siRNA maintenance, we constructed small RNA libraries from control and ds*AGO2 B. germanica*-treated individuals and analyzed the siRNAs produced from a dsRNA for Polyh injected, in a second treatment, to both control and treated individuals. The analysis of the reads produced from heterologous dsRNA allowed us to study the activity of the RNAi machinery, regardless of whether the dsRNA had target mRNA or not. The most abundant length for *B. germanica* siRNAs is 22 nt, a length included in the range assigned to siRNAs [27], but larger than that seen in *D. melanogaster*, where the most abundant size for siRNAs is 21 nt [28,29]. The model these authors propose for determining the siRNA length involves the fact that the phosphate-binding pocket in the fruit fly Dcr-2 binds the 5’monophosphate of the dsRNA. Then, the C-terminal dsRNA-binding domain (CdsRBD) of Dcr-2 binds to the proximal region of the dsRNA substrate, aligning the dsRNA with the RNase III active site in the RNase III domain [29]. Thus, the distance between the phosphate-binding pocket and the RNase III active site should determine the siRNA length corresponding to 22 nt in the case of *B. germanica*.

Furthermore, we observed a certain prevalence of cytosine as the first nucleotide in the case of the 22 nt siRNAs, matching the dsRNA Polyh sequence in the control libraries, but which seems to be lost in *AGO2*-depleted libraries. This points to a slight preference for loading siRNA duplexes in the RISC where cytosine is favored as the first nucleotide in the guide strand. Additionally, in *D. melanogaster*, AGO2-loaded sequences often start with cytosine, both in the case of exo-siRNAs and miR* [22,30].

The siRNAs produced in *B. germanica* after being treated with a dsRNA do not map homogeneously along the corresponding dsRNA sequences, but instead accumulate in certain areas of the two dsRNA strands. The fact that this occurs for heterologous dsRNA, whose siRNAs have no mRNA targets inside the cells, suggests that the dsRNA cleavage and siRNAs selection is independent of the siRNA activity as a guide for RNA degradation. The heterogeneous distribution of the siRNAs along the dsRNA sequence was also observed for the dsRNA corresponding to *AGO2* and *Dcr-1*, which revealed different siRNA patterns which were also different from that of dsRNA Polyh. These patterns differences between sequences and strands are probably related to the dsRNA sequence used in the study, and point to the difficulty of predicting a priori the siRNAs that are produced and maintained.

In the case of dsRNA Polyh, the profiles of the accumulated reads observed in the control and ds*AGO2* animals were similar, but not identical. In the libraries from ds*AGO2* animals, more 22 nt reads were detected, and some of these mapped in areas of the sequence where far fewer reads were found in control animals. The higher proportion of 22 nt siRNAs and their increase in certain areas of the dsRNA sequence in ds*AGO2* libraries indicate that, also in *B. germanica*, AGO2 has a ribonuclease activity that is reduced in the case of ds*AGO2* animals. In addition, the increase of 22nt siRNAs and the decrease of larger reads suggests AGO2 acts on Dcr-2 activity regulation.

One important step in RNAi is the selection of the guide strand from the siRNA duplex released after Dcr-2 digestion. In *D. melanogaster*, R2D2 participates in this process before AGO2 within the RISC takes over the duplex. R2D2 forms a heterodimer with Dcr-2 in such a way that R2D2 binds the siRNA duplex at the extreme with the greatest double-stranded character in terms of bond energy, which defines the orientation of the siRNA duplex when loaded into the RISC [7]. This orientation determines the guide strand as the one presenting the least thermodynamically stable base-pairing at its 5’ end [7,22]. In the analysis of our libraries, we found that in 62 % of the selected duplexes with a high energy difference between the nucleotide bonds at the two extremes (see Material and Methods and supplementary data), the strand with the highest number of reads is the one with the least thermodynamically stable base-pairing at its 5’ end, as in the proposed model. However, in the remaining cases, a considerably high proportion, the other strand is the one with the highest number of reads. For this reason, although it is possibly the case, we cannot fully assert that the model proposed for *D. melanogaster* is fulfilled in *B. germanica*, and we cannot discard the possibility that in this species, and perhaps in other insects, the guide strand selection mechanism is not the same as in *D. melanogaster*, a point that should be studied in more detail.

According to our results and those provided in the literature involving studies of other species, particularly *D. melanogaste*r, we propose that in *B. germanica*, when the cells detect a dsRNA, there is a rapid and transient increase in the expression of *Dcr-2* and *AGO2* triggered by a yet unknown mechanism that senses the dsRNA and activates RNAi machinery. *AGO2* expression is activated by the presence of a dsRNA in the cell, and not by Dcr-2 or the siRNA duplexes produced by Dcr-2 activity, which are the substrate for AGO2 activity. Dcr-2 then cleaves the dsRNA producing 22 nt siRNA duplexes. Dcr-2, in a separate process to dsRNA cleavage [31,32], and together with the double-stranded RNA binding protein R2D2, loads the siRNA duplex into the RISC [7,33].

Although we could not identify the mechanism determining guide strand selection, we did observe a clear preference for some strands over others. We cannot rule out the possibility that guide strands are selected according to thermodynamic criteria [7,22], but we did detect a slight preference for cytosine as the first nucleotide of the guide strand. When depleting *AGO2* through RNAi (ds*AGO2*), more 22 nt siRNAs, corresponding to dsRNA areas that are less well-represented in the case of the controls, were accumulated due to the reduction of the AGO2 RNase activity.

Upcoming studies that increase our knowledge of RNAi machinery and mechanisms will shed even more light on this proceses that is extremely useful for reverse genetics studies in laboratories, as well as for the design of more effective, specific, and eco-friendly pest-control strategies.

## Funding

This work was supported by Spanish Ministry of Economy and Competitiveness and FEDER (grant number CGL2016-76011-R), Secretaria d’Universitats i Recerca, Catalan Goverment (grant number 2017 SGR 1030), Agencia Estatal de Investigación (grant number PID2019-104483GB-I00 / AEI / 10.13039/501100011033), and Agencia Estatal Consejo Superior de Investigaciones Científicas (grant numbers 2019AEP028 and 2019AEP029).

## Notes

### Competing Interest Statement

The authors have declared no competing interest.

