## Supplemental Figure 1 for "siRNA enrichment in Argonaute 2-depleted *Blattella germanica*"

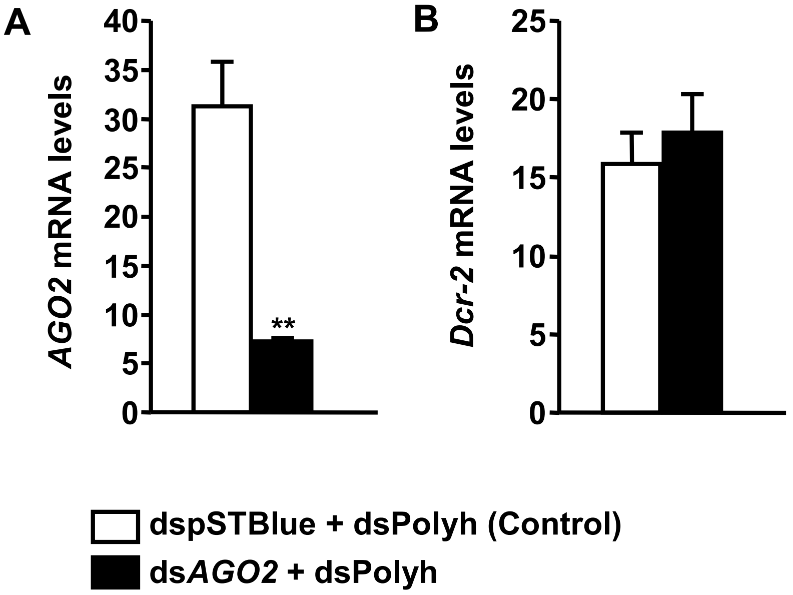


**Figure S1. Depletion of *AGO*2 in samples used to prepare the small RNA libraries**. Freshly emerged penultimate instar nymph (N5D0) were treated with 2 µg of ds*AGO2* or 2 µg of a heterologous dsRNA prepared using using the pSTBlue-1 (dspSTBlue). Five days later (N5D5), both ds*AGO2* and dspSTBlue individuals were treated with 2 µg of ds*Polyh*. Specimens were dissected three days later, in the second day of the last nymphal instar (N6D2), and levels of *AGO2* (A) and *Dcr-2* (B) mRNAs were measured. Data represent copies of mRNA per 1000 copies of *actin-5c* and are expressed as the mean ± S.E.M. (n = 3). Asterisks indicate statistical differences between Control and ds*AGO2* individuals (Student's *t*-test, $^{**}p < 0.01$).
