## Supplemental Figure 2 for "siRNA enrichment in Argonaute 2-depleted *Blattella germanica*"

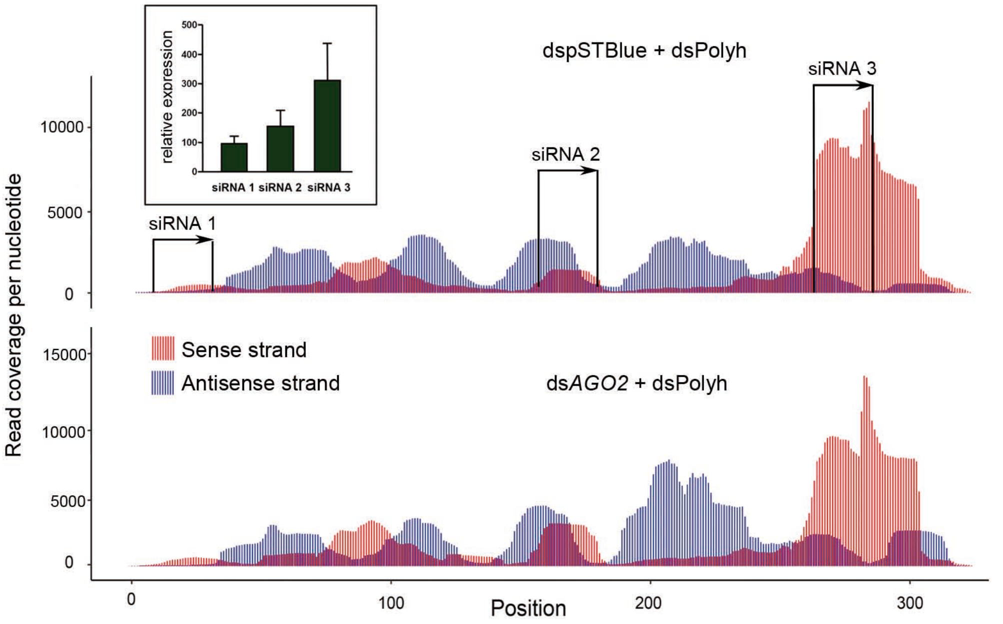


**Figure S2. Analysis of the reads corresponding to dsRNA Polyh in Control and *AGO2*-depleted libraries and checking by qPCR.** Freshly emerged penultimate instar nymph (N5D0) were treated with 2 µg of ds*AGO2* or 2 µg of a heterologous dsRNA prepared using the pSTBlue-1 (dspSTBlue; Control). Five days later (N5D5), both ds*AGO2* and dspSTBlue individuals were treated with 2 µg of a second heterologous dsRNA (dsPolyh). To evaluate the nucleotide coverage (by strand) of the dsRNA Polyh sequence, all the reads between 17 and 50 nt from the three replicates of each treatment, were combined. The inset shows the qPCR quantification of the siRNAs in the Control libraries using as a forward primer the sequences in the position indicated with arrows, and as a reverse primer the SR RT Primer for Illumina provided by the library kit. Y-axis indicates
