## Supplemental data for "siRNA enrichment in Argonaute 2-depleted *Blattella germanica*"

**Supplementary data. Analysis of the reads corresponding to siRNAs duplexes**

It has been proposed that the strand showing the less thermodynamically stable base-pairing at its 5' end, that is, the highest bond energy, is selected as the guide strand (Tomari et al (2004) Science 306, 1377-80; Czech et al (2009) Mol Cell 26,445-56), and then, it will show a higher number of reads than its complementary strand (the passenger strand). This is the model that we will check in our sequences.

In the case of the siRNAs corresponding to the dsRNA Polyh sequence we selected the 15 siRNA showing the highest reads numbers. In the case of dsRNA *AGO2* and dsRNA *Dcr-1*, where we have less reads, we chose the 10 and 9 sequences, respectively, with the higher number of reads. We then matched each sequence with the sequence that would correspond to it after processing by the RNase III Dcr-2 (two nt overhanging at each 3 'end), whether is present in our libraries or not. The energies of the bonds corresponding to the 4 nt at each 5' end were calculated using the ViennaRNA Web Services. Then, the energies of both extremes were subtracted.

Example

**5'-CGUGAUUAGGAUCGUCGAGCCU-3'** 2007 reads
3'-CUGCACUAAUCCUAGCAGCUCG-5' 0 reads

Energy = -3.18
Energy = -4.47 Δ Energies = -3.18 – (-4.47) = +1.29. The results is positive, so, among the pair of siRNAs in the duplex, the one having more reads (the upper one) shows the lowest energy in its 5' end. In this case, the sorting fulfils the proposed model.

We only will apply the model when having Δ Energies > +0.5 or < -0.5 Kcal/mol. Only the strand showing the higher reads number is shown in the tables. The asterisk indicates that that sequence is shown both in the dsPSTBlue + dsPolyh and ds*AGO2* + dsPolyh results. The position is indicated with respect to the sense strand.

dsRNA Polyh in dsPSTBlue + dsPolyh

| siRNA duplex | position | higher vs. lower reads number | Δ Energies (Kcal/mol) | Does it fulfill the proposed model? |
|---|---|---|---|---|
| **CGUGAUUAGGAUCGUCGAGCCU** | 282-303* | 2007 vs. 0 | -3.18-(-4.47) = +1.29 | YES |
| **CCGACUAUGUACCUCAUGACGU** | 263-284* | 1618 vs. 0 | -4.07-(-2.65) = -1.42 | NO |
| **CGACUAUGUACCUCAUGACGUG** | 264-285 | 681 vs. 20 | 0 | - |
| **UUGCGACCCCGACUAUGUACCU** | 255-276* | 607 vs. 23 | -0.15 | - |
| **UUCUUGGUCGUUAACAAUGGGG** | 171-150 | 357 vs. 2 | +1.72 | YES |
| **UGCGUUGCGACCCCGACUAUGU** | 251-272 | 347 vs. 13 | -3.51 | NO |
| **UCGUGUCGGGUUUAACAUUACG** | 76-55 | 278 vs. 7 | -3.85 | NO |
| **CUAUGUACCUCAUGACGUGAUU** | 267-288* | 278 vs. 1 | +2.30 | YES |
| **UGGGUCUAGUGGGACGCAUGUU** | 217-196 | 245 vs. 19 | -2.91 | NO |
| **UUCCUGGCCCAACACGCUCUGC** | 232-253 | 242 vs. 18 | +0.01 | - |
| **UUUCCCUGUAGAACUCUUUUCC** | 121-100* | 255 vs. 0 | -2.16 | NO |
| **UACGGAUUUCCUUGAAGAGAGU** | 58-37 | 20 vs. 1 | +1.10 | YES |
| **UUCCCUGUAGAACUCUUUUCCU** | 120-99 | 211 vs. 0 | -2.49 | NO |
| **CCAGGAAUUUGUAACAACGGUU** | 238-217* | 210 vs. 2 | -0.15 | - |
| **UCAUGAGGUACAUAGUCGGGGU** | 281-260 | 198 vs. 126 | +4.58 | YES |

dsRNA Polyh in ds*AGO2* + dsPolyh

| siRNA duplex | position | higher vs. lower reads number | Δ Energies (Kcal/mol) | Does it fulfill the proposed model? |
|---|---|---|---|---|
| **CGUGAUUAGGAUCGUCGAGCCU** | 282-303* | 4954 vs. 2 | +1.29 | YES |
| **UAGUGGGACGCAUGUUGACAAC** | 211-190 | 2815 vs. 44 | +1.75 | YES |
| **ACGUGAUUAGGAUCGUCGAGCC** | 281-302 | 2583 vs. 13 | +1.24 | YES |
| **CCGACUAUGUACCUCAUGACGU** | 263-284* | 1962 vs. 16 | -1.42 | NO |
| **UCUUGGUCGUUAACAAUGGGGA** | 170-149 | 1598 vs. 1 | +3.96 | YES |
| **UCUAGUGGGACGCAUGUUGACA** | 213-192 | 1555 vs. 147 | +0.41 | - |
| **UUGCGACCCCGACUAUGUACCU** | 255-276* | 1213 vs. 165 | -0.15 | - |
| **CUAUGUACCUCAUGACGUGAUU** | 267-288* | 1203 vs. 14 | +2.30 | YES |
| **UAACGACCAAGAAGUGAUGGAU** | 159-180 | 1192 vs. 38 | +2.21 | YES |
| **CCCGACUAUGUACCUCAUGACG** | 262-283 | 1083 vs. 489 | -4.58 | NO |
| **CAACGGUUGGGUCUAGUGGGAC** | 224-203 | 997 vs. 51 | +3.04 | YES |
| **AGGAAUUUGUAACAACGGUUGG** | 236-215 | 899 vs. 47 | -0.90 | NO |
| **CCAGGAAUUUGUAACAACGGUU** | 238-217* | 895 vs. 41 | -0.15 | - |
| **UUUCCCUGUAGAACUCUUUUCC** | 121-100* | 876 vs. 10 | -2.16 | NO |
| **CGACUAUGUACCUCAUGACGUG** | 264-285 | 858 vs. 166 | 0 | - |

dsRNA *AGO2* in ds*AGO2* + dsPolyh

| siRNA duplex | position | upper sequence vs lower sequence reads | Δ Energies | Does it fulfill the proposed model? |
|---|---|---|---|---|
| **UCCGAUCGUUCAGACAUAGAAU** | 240-219 | 377 vs. 85 | -2.95 | NO |
| **UUCAGACAUAGAAUAAUCCAUU** | 232-211 | 266 vs. 7 | +2.49 | YES |
| **UAGAAGAAUUUCUGAGGACUAU** | 367-388 | 253 vs. 0 | +1.53 | YES |
| **UUGAACGGAUUGGUGGACGAGU** | 79-100 | 192 vs. 0 | +1.67 | YES |
| **AAUGAGAGGCGUGUGGAAUGCA** | 158-179 | 133 vs. 0 | 0 | - |
| **AUUCUAUGUCUGAACGAUCGGA** | 219-240 | 133 vs. 0 | +1.55 | YES |
| **CGUCCGAUCGUUCAGACAUAGA** | 242-221 | 122 vs. 0 | -3.17 | NO |
| **UGAGAGGCGUGUGGAAUGCAAA** | 160-181 | 115 vs. 6 | +0.46 | - |
| **AAUAGAAGAAUUUCUGAGGACU** | 365-386 | 108 vs. 0 | +4.67 | YES |
| **CACUGCUCGCAUAGAACUCCUU** | 201-180 | 105 vs. 0 | +2.03 | YES |

dsRNA *Dcr-1* (from Ylla et al., 2016)0

| siRNA duplex | position | upper sequence vs lower sequence reads | Δ Energies | Does it fulfill the proposed model? |
|---|---|---|---|---|
| **AAUUUGAACAGAUGUAGUACUG** | 130-109 | 1183 vs. 0 | +4.04 | YES |
| **AUGGCUAGGAAUCUGUGUUCUU** | 25-46 | 837 vs. 0 | -0.46 | NO |
| **AGAUGUAGUACUGCCAACAGGU** | 121-100 | 342 vs. 259 | +1.35 | YES |
| **AAGAGACUGUAGUAAGUAGCUU** | 328-307 | 236 vs. 0 | +2.01 | YES |
| **AAGUUUCUGUAUUAUCACCGUU** | 152-131 | 204 vs. 10 | +3.36 | YES |
| **UGAGUAAGAGACUGUAGUAAGU** | 333-312 | 200 vs. 18 | -2.31 | NO |
| **GACUGUAGUAAGUAGCUUCUGU** | 324-303 | 199 vs. 0 | -1.94 | NO |
| **CGUUAAUUUGAACAGAUGUAGU** | 134-113 | 181 vs. 0 | -0.46 | - |
| **AUUUGAACAGAUGUAGUACUGC** | 129-108 | 179 vs. 18 | +2.78 | YES |
