## Supplemental Table 1 for "siRNA enrichment in Argonaute 2-depleted *Blattella germanica*"

Table S1: Reads in small RNA libraries from N6D2 females treated with dspSTBlue + dsPolyh and ds*AGO2* + dsPolyh

| Library | Raw reads | Mapped reads to *B. germanica* genome | Mapped reads to dsRNA Polyh | Mapped reads to dsRNA *AGO2* | Total maped reads | % of Polyh mapped reads respect all the reads mapped to the genome and to Polyh sequence | % of *AGO2* mapped reads respect all the reads mapped to the genome including the Polyh sequence |
|---|---|---|---|---|---|---|---|
| dspSTBlue + dsPolyh_1 | 3845773 | 2274577 | 25052 | 5 | 2299629 | 1,089 | 0,000217426 |
| dspSTBlue + dsPolyh_2 | 4796432 | 2517076 | 37549 | 4 | 2554625 | 1,470 | 0,000156579 |
| dspSTBlue + dsPolyh_3 | 105164 | 61884 | 395 | 5 | 62279 | 0,634 | 0,008028388 |
| ds*AGO2* + dsPolyh_1 | 3914306 | 2086285 | 58335 | 3952 | 2144620 | 2,720 | 0,18427507 |
| ds*AGO2* + dsPolyh_2 | 4681950 | 2689828 | 58973 | 5515 | 2748801 | 2,145 | 0,200632931 |
| ds*AGO2* + dsPolyh_3 | 3567950 | 1799842 | 56413 | 13674 | 1856255 | 3,039 | 0,73664448 |
