## Supplemental Table 2 for "siRNA enrichment in Argonaute 2-depleted *Blattella germanica*"

Table S2: Reads in small RNA libraries from N6D2 females treated with dsPolyh and ds*Dcr-1* (data from Ylla et al., 2016)

| Library | Raw reads | Mapped reads to *B. germanica* genome | Mapped reads to dsRNA Polyh | Mapped reads to dsRNA *Dcr-1* | Total maped reads | % of Polyh mapped reads respect all the reads mapped to the genome and to Polyh sequence | % of *Dcr-1* mapped reads respect all the reads mapped to the genome including the Polyh sequence |
|---|---|---|---|---|---|---|---|
| dsPolyh_1 | 5319915 | 3242301 | 8982 | 52 | 3251283 | 0,2770 | 0,0016 |
| dsPolyh_2 | 4324534 | 2994466 | 15301 | 133 | 3009767 | 0,5110 | 0,0044 |
| ds*Dicer-1* _1 | 5689294 | 3557940 | | 21854 | 3557940 | | 0,6142 |
| ds*Dicer-1* _2 | 4520535 | 2797709 | | 16811 | 2797709 | | 0,6009 |
